# The Clinical Kinase Index: Prioritizing Understudied Kinases as Targets for the Treatment of Cancer

**DOI:** 10.1101/2020.02.14.943886

**Authors:** Derek Essegian, Rimpi Khurana, Vasileios Stathias, Stephan C. Schürer

## Abstract

The approval of the first kinase inhibitor, Gleevec, in 2001, ushered in a paradigm shift for oncological treatment—the use of genomic data for targeted, efficacious therapies. Since then, over 48 additional small molecule kinase inhibitors have been approved, solidifying the case for kinases as a highly druggable and attractive target class. Despite the established role deregulated kinase activity plays in cancer, only 8% of the entire kinome has been effectively “drugged”. Moreover, a quarter of the 634 human kinases are vastly understudied. We have developed a comprehensive scoring system which utilizes differential gene expression, clinical and pathological parameters, overall survival and mutational hotspot analysis to rank and prioritize clinically-relevant kinase targets across 17 solid tumor cancers from The Cancer Genome Atlas (TCGA). Collectively, we report that dark kinases have potential clinical value as biomarkers or as new drug targets that warrant further study.

## Introduction

The human genome encodes about 634 kinases (pseudokinases included). However, as of 2019[1], only 49 primary kinase targets (∼8%) are currently targeted by FDA approved oncology drugs. Furthermore, 70% of these approved cancer kinase drug targets belong to the TK group (Tyrosine Kinase). For several cancers though, targeting the TK group has not been an effective strategy, despite overwhelming evidence of kinase dysregulation in those tumors [2]. For example, TKIs (Tyrosine Kinase Inhibitors) have shown little to no clinical efficacy in the treatment of bladder, esophageal, prostate, brain and stomach cancer [3] [2, 4-6]. While it has been firmly established that aberrant kinase activity indeed leads to cancer progression and metastasis in these cancers, researchers have not identified ideal cancer-specific kinase targets or novel drug combinations to improve standard of care.

With over 175 kinase drugs currently in clinical trials (www.icoa.fr/pkidb/), newer targets are also being evaluated including AKT, Aurora Kinases, CHEK1 and CDK1. Despite the large number of drugs that are being investigated, the majority are for well-known, previously approved kinase targets such as EGFR, VEGFR, PI3K and mTOR. Nevertheless, there are no small molecule drugs that target kinases in the CAMK, CK1 or AGC group of kinases as their primary target (Ki < 10 nM), despite increased evidence for their clinical relevance in cancer [7, 8]. The extent of such a misrepresentation has been highlighted by the Illuminating the Druggable Genome (IDG, https://druggablegenome.net/) project, where 23.8% (151) of the 634 kinases are labeled as “Understudied” as they lack GeneRIFs, antibodies, citations in the literature and potent chemical probes [9]. Consistent with this is the fact that the current kinase inhibitors target not only a narrow range of targets, but also a narrow range of pathways including Angiogenesis, Cell Adhesion, Immune System Signaling (Cytokine, TCR, BCR) and anti-apoptotic pathways[10]. For example, all kinase inhibitors for renal cell carcinoma target angiogenic pathways [11]. It is probable that there exists a strategy for targeting multiple, orthogonal pathways that work in a synergistic manner, as opposed to targeting kinases with overlapping pathways. The discovery of novel kinase targets may lead to synergy efforts to target orthogonal signaling pathways which will be a crucial strategy for treating the complex disease of cancer [12]. It has already been shown that this strategy may prevent or reduce the incidence of resistance pathways and kinome reprogramming, which is inevitable upon singular treatment with a highly specific kinase drug such as EGFR inhibitor, lapatinib [13, 14].

The typical 20-year long, multi-billion dollar drug discovery pipeline all begins with target identification and prioritization [15]. This is arguably one of the most important steps as drug failure in the clinic is often due to lack of efficacy or due to toxicities from poor target choice [15, 16]. The use of large-scale -omics data has streamlined this process by allowing researchers to combine multiple parameters to evaluate a protein’s potential as a drug target or biomarker. Often the first glimmer of target potential arises from the analysis of RNA and protein expression in disease tissue compared to normal, healthy tissue. The Cancer Genome Atlas (TCGA) is one such publicly available database where a breadth of information (transcriptomic, genomic, proteomic, clinical, pathological and histological) is available for this investigation. An ideal drug target, as described by Bayer [17], is one that is first and foremost, “druggable” and “assayable”-which all kinases typically are. This target will also have an activity that is disease specific; i.e. it is differentially expressed and active in diseased tissue compared to normal tissue. This can be determined preliminarily by differential gene expression analysis and proteomic or phosphoproteomic studies. The target should not be uniformly distributed and expressed throughout the body; a characteristic that can be checked using expression databases such as GTex [18] or The Human Protein Atlas[19]. Also, a favorable IP situation is more likely for “dark” and understudied kinases where there are few or no known small molecule inhibitors. Finally, and the most difficult parameter to satisfy using informatics alone, kinase targets have to be disease modifying or have a proven function in the pathophysiology of disease. While dysregulation of the kinome as a whole has been indicated in the initiation and progression of nearly every cancer type [20], disease modifying kinases must be identified per cancer type using a combination of genetic perturbations and biochemical analyses. Datasets such as DepMap [21] may serve to be an important resource to help prioritize and rank order drug targets for “disease modifying” potential before pursuing further target validation studies. In totality, combining multiple large-scale multi-omics datasets can be a useful first step in prioritizing novel kinase target lists prior to conducting any additional cell-based or animal-based experiments.

We herein describe a kinase target prioritization index using data from The Cancer Genome Atlas (TCGA) across 17 cancers. By combining Differential Gene Expression (DGE), Kaplan-Meier survival, mutational hot-spot and clinical/pathological correlation analyses, we have developed a scoring system to rank-order *clinically* relevant kinase targets for each cancer cohort (Figure 1). In short, we have analyzed and prioritized kinases whose mRNA expression levels appear to be prognostic and correlate positively with progression of cancer through TNM staging, histological grading and overall survival (OS). Since kinase activity does not always correspond with mRNA expression levels (e.g. mTOR oncogenic activity is due to increase in phosphorylation)[22], we also leveraged TCGA genomic data and prioritized kinases that confer a selective advantage to tumor development as measured by the accrual and clustering of mutations at specific regions of their amino acid sequence. Moreover, by integrating with external target annotation resources, we evaluated the kinase index scores based on a number of clinically relevant classifications such as target development level (TDL), which classifies a target based on available target validation knowledge, kinase family class (which corresponds to phylogenecity and substrate), and MOA (Mechanism of Action) of approved drugs for each cancer-type [9, 23].

**Figure 1:**
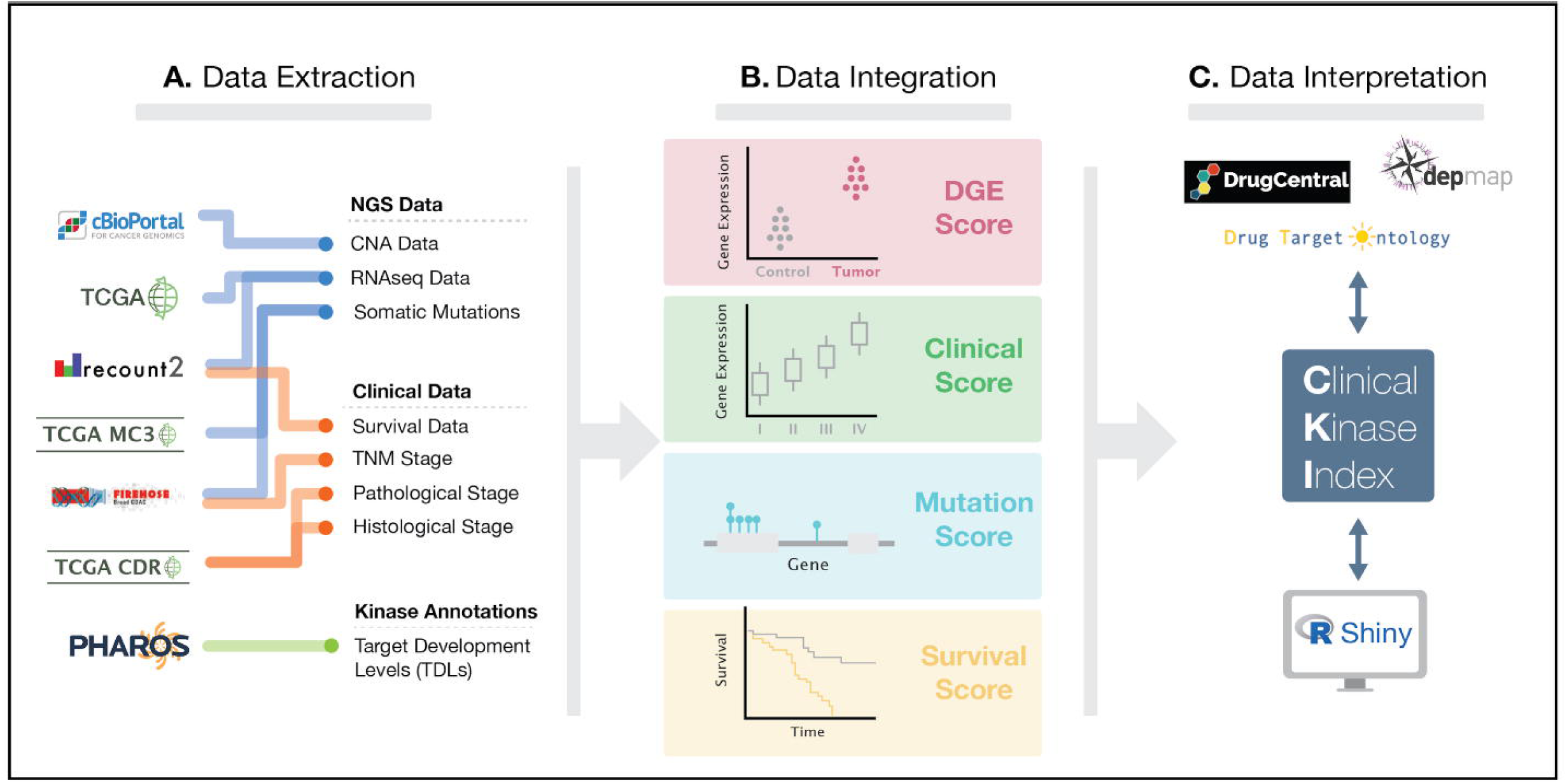
Overview of workflow A.) Data was extracted from multiple sources for curation, filtering and normalization. B.) mRNA levels and mutational hotspots were analyzed to generate differential gene expression analyses and mutational frequencies. mRNA levels were also correlated with survival and clinical/pathological outcomes. Various statistical methods were employed and all significant correlations were scored. C.) Final Clinical Kinase Index (CKI) scores were generated and mapped to other data for further analysis and interpretation.

We have developed an App to explore the clinical relevance of all kinases. All scoring, classifications, normalized data and statistical analyses are available via the Clinical Kinome Index (CKI) App at http://cki.ccs.miami.edu. To use the app, one can simply select the “Gene” tab and choose a kinase of interest. The CKI scores for each cancer for this kinase will be generated in table form, along with target development level, rank, kinase group and family and whether or not this kinase has an approved drug MOA. Many annotations from the Drug Target Ontology (DTO) [11] are available as facets to filter and select kinase targets. To start from a disease of interest, one can select a TCGA cancer in the “Disease” tab. A volcano plot will be generated where all differentially expressed genes are displayed. One may click points on the plot to see the specific gene, its count-per-million and log fold change (compared to normal tissue). If the “Study” sub-tab is selected, cancer and gene may be chosen to display box-plots representing mRNA levels for each T, N and M stage. Finally, if “Survival” sub-tab is selected, a Kaplan-Meier plot is generated for the cancer and gene pair of interest. All data tables may be downloaded via the Download Data tab.

Overall, our study provides a valuable resource for the scientific community where the clinical relevance of kinase genes across solid-tumor cancers can quickly be evaluated, especially for understudied kinases and cancers for which no approved first-line kinase therapy exists.

## Results

### Understudied calcium/calmodulin-dependent kinases are highly overexpressed across multiple cancer types

The NIH Illuminating the Druggable Genome Project (IDG) research consortium (https://druggablegenome.net/) has curated a list of “Understudied Kinases”, which were selected using a combination of bibliometric and other measures including lack of R01 funding, limited GeneRIF and GO annotation, and lack of available potent and specific chemical probes. These include kinases that bear the target development levels of Tchem, Tbio and Tdark. Despite being largely ignored by the scientific community, understudied kinases are gaining more attention due to their novelty and therefore opportunities and clinical importance [7, 8]. Of the 151 “understudied” kinases, 22% are from the “Other Group” (which include the subfamilies of *BUB, AUR* and *PLK*), 20.7% are of the CGMC group (which contain MAP-kinases and Cyclin-dependent kinases) and 17.3% are CAM Kinases. The rest of the understudied kinases include AGC and Atypical kinases, with the lowest number of kinases belonging to the already very well-studied TK and TKL groups (Figure 2A, 2B).

**Figure 2:**
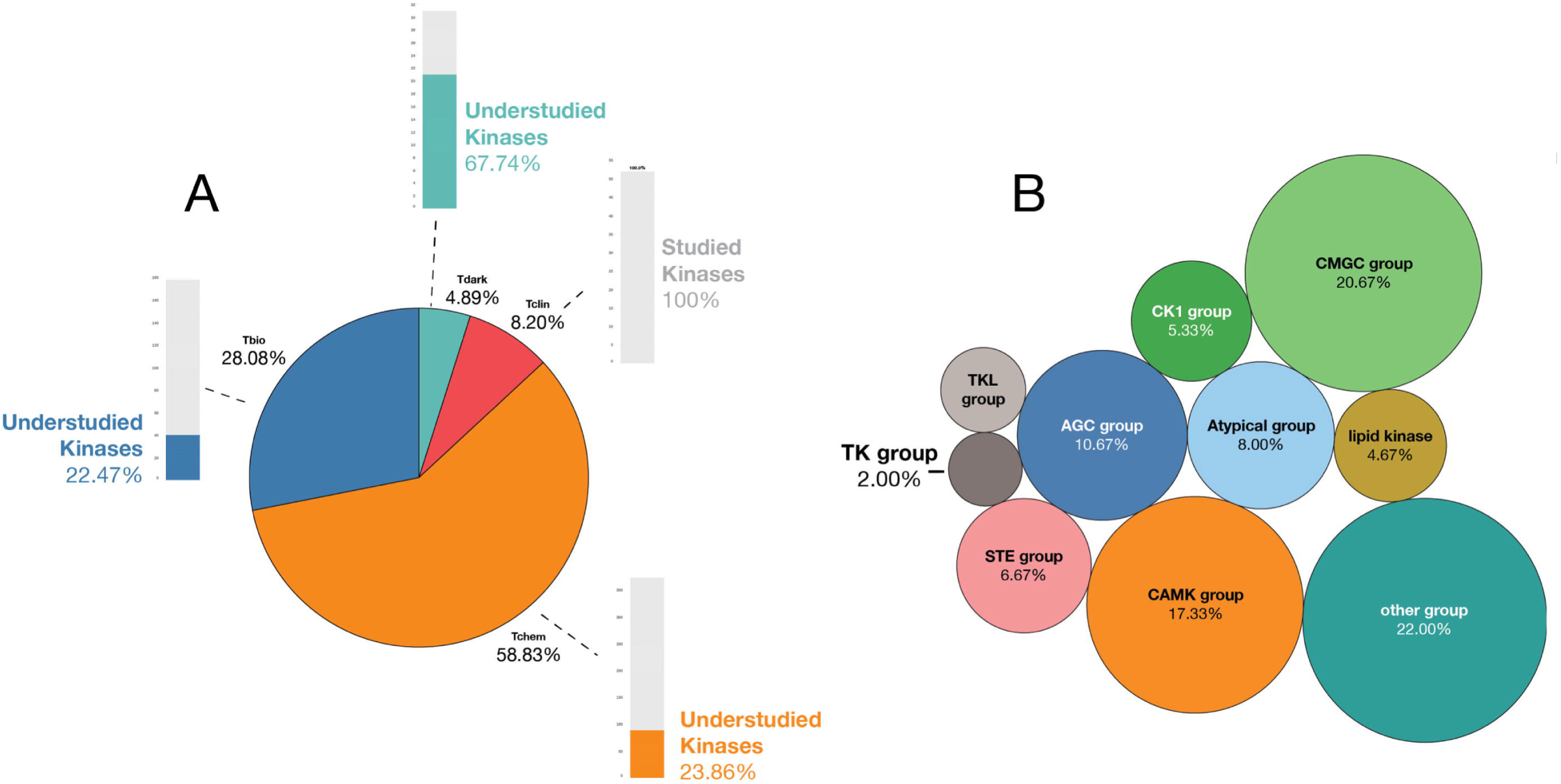
Analysis of targets used in this study. A.) There are 634 kinases annotated on Pharos. 8.2% are considered Tclin. 151/634 are considered ‘‘Understudied’’; these represent kinases that bear Tdark, Tbio and Tchem annotations. B.) Certain groups of Kinases are historically understudied include those from the ‘‘CMGC’’ group, ‘‘CAMK’’ group and ‘‘Other’’ group.

Since differential gene expression analysis is one of the most effective, scalable and predictive methods for target prioritization, we proceeded with a comprehensive analysis of the expression patterns of each of the 624 kinases across 19 TCGA cancer types. Kinases are the signaling molecules for a variety of cellular functions often hijacked by cancer. Therefore, it is not surprising that 424 of the 624 kinases studied were found to be significantly differentially overexpressed (p-value <0.05) in at least two cancers in the TCGA (Figure 3A, Supplemental Table 1). *BUB1, BUB1B* and *PLK1* are all significantly overexpressed in every cancer analyzed in this study. In addition to being commonly differentially overexpressed, the average Log_2_FC between normal and tumor cells is >2 for these kinases. Interestingly, these three kinases all interact with one another in the kinetochore-microtubule spindle assembly checkpoint during mitosis [24, 25]. Several other kinases involved in mitosis/cell cycle are also overexpressed in the majority of cancers (*AURKA, AURKB, CDK1*) [26-28], underscoring a hallmark of cancer; cell cycle dysregulation. Many campaigns are well underway in the pre-clinical and early clinical trial phases to assess the efficacy of targeting such cell cycle kinases; some with great potential [29, 30].

**Figure 3:**
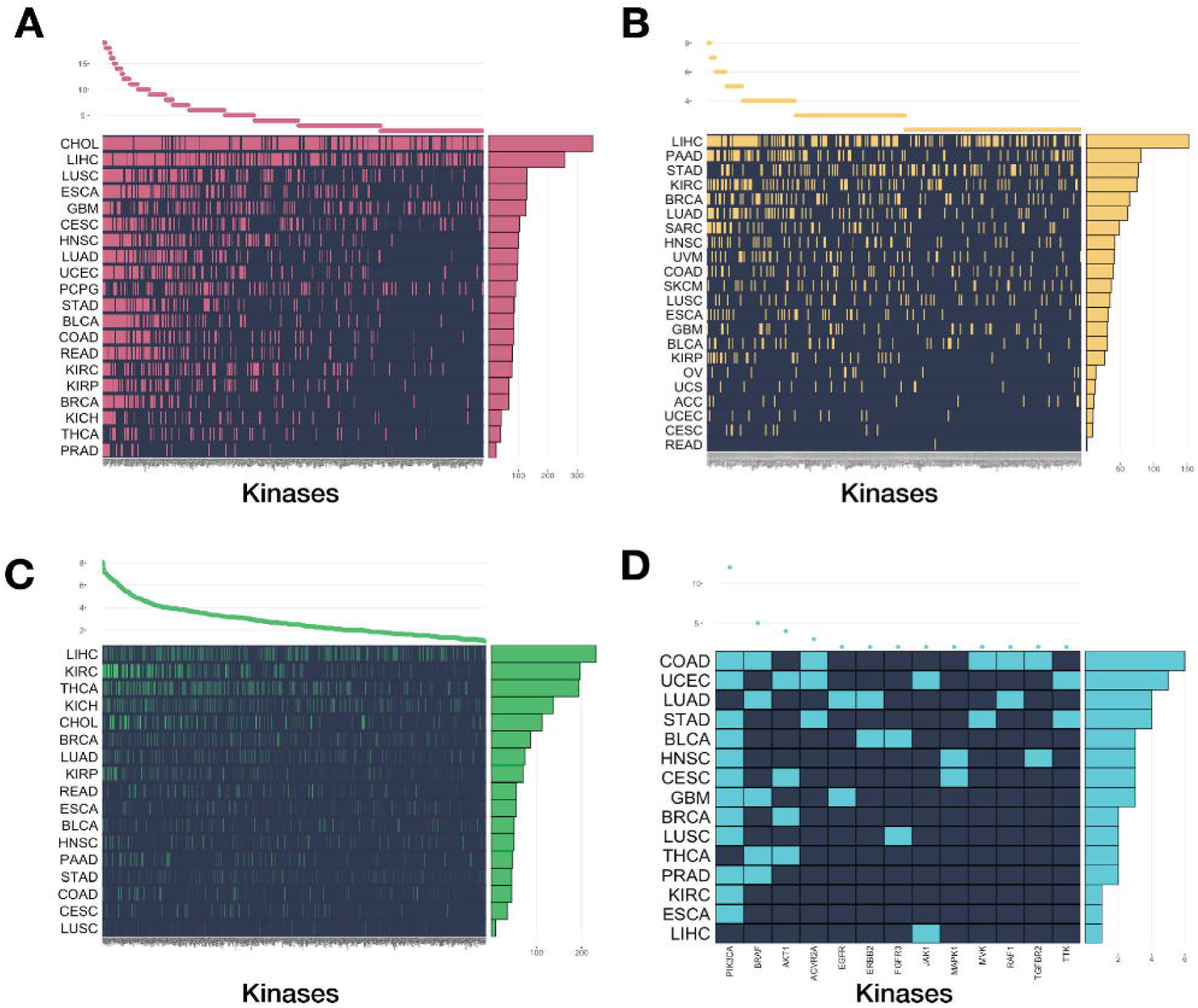
4 Analyses which contributed to the Clinical Kinase Index (CKI). The X-axis contains all kinase names in the analysis while the Y-axis represents the TCGA cancer. The plot on the top of the graph shows how many cancers a particular kinase is “significant” in. The plot on the right side of the graph represents how many significant kinases a particular cancer has for this analysis. A.) 424 kinases are differentially overexpressed in at-least 2 cancers. B.) 317 kinases are prognostic of survival in at least 2 cancers. C) Many Kinases have expressions which correlate with T, N and M staging. This heat-map quantifies the extent to which each kinase correlates with the clinical outcome, TNM Staging (Total score out of 3, Max score of 1 per each parameter) D.) 13 kinases have significant hotspot mutations in at least 2 cancers.

The differential gene expression analysis included 149 understudied kinases across 19 cancers from the TCGA. 102 understudied kinases were shown to be differentially overexpressed in at-least 2 cancers (Figure S1). In every cancer cohort, the ratio of overexpressed understudied kinases compared to well-studied kinases was nearly 1:1.

Several kinase groups appear to be enriched in their overexpression in various cancers. Averaging the log_2_FC of all kinases across all cancers demonstrates that certain kinase transcripts are consistently very highly upregulated. Kinases from the historically understudied CAMK group have the highest average log_2_FC (2.14) (Supplementary Figure 2). 24 of the 26 understudied CAMK kinases are overexpressed in at least one cancer. The most commonly overexpressed CAMK is the pseudokinase *CAMKV*, which is significantly upregulated in 14 cancer cohorts. The understudied multi-functional CAMK kinases (*CAMK1D, CAMK1G, CAMKK1, PNCK*) are overexpressed in 12 cancers, with *PNCK* being overexpressed in 9 alone. *PNCK* (Tbio) mRNA expression and activity has recently been linked to renal cell carcinoma progression and survival [10], breast cancer tumor microenvironment remodeling [31] and decreased sensitivity to chemotherapies such as temozolomide [32]. *PNCK* is highly overexpressed in KIRC (>5 log_2_FC), LUSC (>6 log_2_FC), and LIHC (>6 log_2_FC). *PNCK*, in fact, is the most significantly overexpressed kinase in these cancers, suggestive of a tumor-specific differential need for *PNCK* or CaMK activity compared to normal tissue. GTex and other human proteomic/transcriptomic studies reveal *PNCK* has very low expression levels in normal adult tissue, with the highest expression of mRNA and protein found in the dentate gyrus of the hippocampus. Non-specific CaM-kinase inhibitor, KN-93, has been shown in pre-clinical cancer cell models to induce cell-cycle arrest and apoptosis [33, 34]. Despite these data, none of the CAMK kinases are currently targeted by FDA drugs nor are they being evaluated in clinical or (published) pre-clinical studies.

### Mutational hotspots found in several understudied kinases may correlate with overall survival

Many studies have demonstrated that somatically acquired mutations in kinase domains lead to tumorigenesis and promote cancer progression [35-37]. As mutations accumulate in a precancerous cell, some mutations confer selective advantage through the promotion of tumorigenic functions (such as unchecked cell cycle progression, immune evasion, invasion and metastasis etc.), whereas others are effectively neutral “passengers” and are the byproduct of “driver” mutations. The discovery of frequent mutations in various kinase active sites has given rise to a new approach in drug development. Selectively targeting the mutated version of the kinase (versus the wild-type version) has led to great clinical success in oncology. For example, *BRAF* V600E inhibitors have greatly improved survival outcomes in melanoma patients with said mutation[36]. Thus, it is important to prioritize kinases with driver mutations as potential novel drug targets. Many cancer genes form mutational hotspots that disrupt their functional domains or active sites, leading to gain- or loss-of-function [38]. Therefore, we performed a mutational hotspot analysis on the entire kinome across 20 cancer types. 42 kinases in total were found to have significant hotspot mutations, with kinases from the TK family being mutated most frequently. The most commonly mutated kinases were *PIK3CA* (12 cancers), *BRAF* (5 cancers) and *AKT1* (4 cancers), all of which have mutant-targeted inhibitors in clinic or in trial (Figure 3D, Supplemental Table 2) [36, 39, 40]. 8 understudied kinases were found to have hotspot mutations in at-least 1 cancer (*CDC42BPA, DYRK1B, DYRK4, LMTK3, MAPK15, NEK7, TSSK1B* and *TTBK1*), with *DYRK4, LMTK3, MAPK15, NEK7* all having significant hotspot mutations in stomach adenocarcinoma (STAD). Additional study showed that *DYRK1B* was mutated in 4.27% of Endometrial Carcinoma samples (UCEC) with worse overall survival for the mutated kinase (Supplementary Figure 3). Further work must be done to determine which of these kinase mutations are driver mutations and which downstream genomic and transcriptomic effects these mutations have on the tumor. STAD and COAD proved to be the cancers with the highest kinase mutational burden with 16 and 11 kinases significantly mutated, respectively. There are currently no first-line treatments for gastric adenocarcinoma that include kinase inhibitors. As a highly heterogeneous disease, genomic data obtained from studies such as this may usher in a new era of personalized medicine for gastric cancer with novel kinase inhibitors against clinically relevant but rarely amplified and mutated kinases.

Comparing our mutational analysis to other pan-cancer TCGA analysis confirms that few kinases are significantly mutated, and few mutations are prognostic of survival. Smith and Sheltzer [41] identified *all* non-silent mutations with >2% frequency and use Cox-proportional hazard analysis to detect genes prognostic of survival. Analysis of their hazard ratios as Z-scores does show that Tdark kinase *NRK* is significantly mutated and prognostic in HNSC. Additionally, understudied kinases *CAMK1D, PNCK* and *NEK3* have mutations with significant pan-cancer prognostic value, with *PNCK* being mutated frequently in LUAD and *NEK3* in COAD/READ. Various large-scale mutational analyses of tumors all confirm that the vast majority of somatically acquired mutations are passenger mutations of little or no functional consequence that arise simply as a result of the random mutagenic processes underlying the development of cancer[42]. It is rare that one single kinase is commonly mutated (with the very well-known exception, BRAF, which is mutated in over 60% of melanoma cases) [42], suggesting that several infrequently mutated kinases most likely contribute to tumorigenesis.

### Copy-number alterations and gene amplifications are frequent among dark kinases

There are 31 “Dark Kinases” (Tdark) for which very little information is known. More specifically, these kinases have less than 50 antibodies in Antibodypedia, less than 3 Gene RIFs and a Jensen Lab PubMed text mining score of less than 5 [23]. Tdark kinases are further characterized by poorly defined roles in wider signaling networks, poorly defined function and regulation, poorly defined kinase substrates, lack of activation-loop phospho-antibodies or immunohistochemistry-grade antibodies, and lack of selective chemical tools for functional characterization. Over a quarter of the dark kinases belong to the “Other” group of kinases. Dark kinases are highly represented amongst the AGC Group (13.3%), Atypical Kinases (10%), and non-protein “Small molecule kinases” (10%). A number of kinases from the so called “Ignorome” *are*, in fact, known to interact with FDA-approved multi-kinase inhibitors. For example, according to DrugCentral data [1, 43], crizotinib, ruxolitinib, nintedanib, vandetanib, bosutinib, sorafenib and sunitinib all inhibit understudied kinases in the low nanomolar range (Including *SBK3, STK32A, RIOK1, CDK15* and *CSNK1A1L*), one of which is Tdark (CSNK1A1L) [1]. Thus, it is likely that some therapeutic effect or anti-cancer phenotype can be achieved through inhibition of dark kinases. Due to the homology of kinase ATP binding sites across the kinome and the nature of many kinase inhibitor chemotypes to target different kinases, it is very likely that Tdark kinases can be effectively targeted with small-molecule inhibitors. To promote the pre-clinical study of these dark targets, we must first evaluate and analyze the available clinical data to prioritize the kinases based off of several criteria including mutational status, frequency of genetic alteration, and differential gene expression compared to normal samples. If these genomic or transcriptomic variations then correlate with clinical or pathological outcomes, the kinases could be explored in-depth as potential oncogenic drivers.

22 of the dark kinases are significantly overexpressed in at least one TCGA cancer cohort. Increased expression of 15 Tdark kinases have been shown to correlate with decreased overall survival across multiple cancers. For example, high ADCK5 mRNA levels are a negative prognostic indicator in renal cell carcinoma, liver hepatocellular carcinoma, uterine carcinoma and uveal melanoma. In breast cancer, overexpression of 6 dark kinases is associated with decreased overall survival (*ALPK3, CSNK1A1L, CSNK2A3, NRK, POMK* and *PSKH1*) (Supplemental Table 2, 3).

We have found that many Tdark kinases have altered genetics in the TCGA dataset. Although no Tdark kinases were detected to have significant hotspot mutations, querying all dark kinases across 9,519 samples in the TCGA (for which CNA/CNV data are available) in 24 cancer cohorts, we found that these genes have altered copy numbers in 36% of patients. *NRBP2, POMK, ADCK5, SCYL3, PSKH2*, and *ETNK2* are amplified in over 5-10% of all patients. While CNA and CNVs do not linearly correlate with increases or decreases in mRNA expression, the potential increased expression of many kinases in primary human tumors and their location in focal amplification peaks with other cancer promoting genetic alterations suggests that dark kinases have important functions for the tumor cell phenotype that have not been characterized to date.

It was expected that Tclin kinases would generally score higher on our clinical kinase index than other less-characterized proteins. In BLCA, CHOL, HSNC, KIRC, LUAD, READ, STAD cohorts, CKI scores were highest for Tclin and MOA-cancer specific targets (TMOA). Surprisingly, in several cancer cohorts (BRCA, PRAD, COAD, LUSC), Tdark kinases scored higher on the CKI scale than known clinical targets (Tclin). Additional analysis of the dark kinome in these cohorts was performed to further prioritize these kinases. Out of 1,108 breast cancer patients, for example, dark kinases are altered in 54% of samples. Overall survival is significantly worse in patients with dark kinase alterations versus those without (p= 4.90E-3). Multiple kinases are commonly amplified including *ADCK5* (14% of samples), *ETNK2* (13% of samples), *NRBP2* (15% of samples), *PSKH2* (12%), *RPS6KC1* (11%) and *SCYL3* (10%). Though, gene amplification of only UCKL1 (6%) and SBK2 (3%) correlated negatively with overall survival and progression free survival in a Kaplan-Meier univariate analysis (p<0.05). Cox-proportional hazard ratios on Tdark kinases and their CNAs reveal that *SBK2, ETNK2, PSKH2, SCYL3*, and *NRBP*2 are all significantly prognostic for survival across all cancers [41]. Genes that are amplified in a peak are likely involved in the driving of oncogenic pathways and alterations. *ETNK2* is significantly focally amplified in breast cancer tumors (Q= 7.6E-5) and located in a peak with 94 other genes[44]. *ADCK5, ETNK2, PSKH2, RPS6KC1* and *SCYL3* are all significantly focally amplified in breast cancer samples as well, but are not located in a peak region. Similar trends were confirmed in prostate, lung and colon cancers (Supplemental Table 2).

### CAMK kinases are potential prognostic biomarkers

The identification and prioritization of novel biomarkers in cancer can be achieved by the integration of gene expression data with clinical data of patient samples. Genes that are prognostic for various cancer typically have expressions that correlate negatively with OS, PFS and tumor progression (TNM staging). Despite the abundance of studies exploring the prognostic value of many genes and kinases, very few prognostic biomarkers exist in clinical practice [45]. FDA approved prognostic biomarkers typically include RNA expression panels of multiple genes. It is often true that good prognostic biomarkers may also be drug target candidates if functional characterization of the gene proves as such [45]. To assess the potential of utilizing kinase expression as prognostic biomarkers, we aggregated for each kinase, RNAseq and clinical data from 17 TCGA cancer types and performed Kaplan-Meier survival analysis with logrank tests and ANOVA tests between the different tumor stages and grades. Kinases were scored separately for significance in each clinical parameter (TNM staging & histological grade when available). A cumulative “clinical score” was generated to rank order kinases that have expression levels which correlate with multiple clinical parameters.

In our analysis, 357 kinases were shown to be prognostic of M-stage, 522 for N-stage and 552 for T-stage. 24 of 31 dark kinases showed correlation with TNM staging in at least one cancer (Figure 3C). The clinical scores were not significantly different between different target development levels or between “studied” and “understudied” kinases. However, average clinical score per kinase phylogenic group showed significant differences in each cancer cohort. The kinases with the highest average clinical scores include *NEK2, TRIB3, MELK, EPHA2, SPEG* and *BRSK1*- several of which are CAMK kinases. *TRIB3* (Tbio), for example, shows correlation with metastasis in 5 cancer cohorts. A member of the CAMK family, literature and other studies confirm that this kinase may play a role in promoting metastasis in lung and colorectal cancers via induction by the transcription factor, NF-kappaB [46-48]. Other CAMK members *CHEK2, TRIB2, STK17A* and *STK17B* are also shown to correlate with TNM staging, further highlighting the potential clinical use for CAM-kinases as novel cancer targets (Supplemental table 4). 317/624 kinases were shown to correlate with survival in at least two cancers (Figure 3B). The kinases with the highest survival scores have been well described in the literature as predictive and prognostic biomarkers in multiple cancers; notably *PGK1, PLK1*, and *AURKA [49-51]*. Less studied kinases such as ALPK3 (Tdark) (Figure S4) and SPEG (Tbio) also were shown to correlate with survival in 6 cancer cohorts (Supplemental table 4).

### The Clinical Kinase Index (CKI) predicts relevance of validated kinase targets

The Clinical Kinase Index (http://cki.ccs.miami.edu/) is the first attempt to rank order and categorize the clinical relevance of the entire kinome, with special focus on understudied kinases (Figure 4). Our CKI scoring system (see Methods) accurately predicts currently in trial or approved drug targets for several cancers in the TCGA. As one might expect, the majority of highly clinically relevant targets would have been or are currently under investigation. For example, breast tumor kinase (*PTK6*) is the highest ranking kinase for BRCA. This kinase is regarded the key regulator in the oncogenic transformation of breast cancer, is overexpressed in >80% of breast tumors and is a highly attractive drug target [52-54]. Other high ranking kinases include those that are already in clinical trial or are FDA approved targets, including PI3K-kinases, MAP-kinases, cyclin dependent kinases and Aurora kinases. In the LUAD cohort, *EGFR* and *MEK1* rank highly; they represent targets of an approved lung cancer drug and compounds under investigation in several clinical trials [35, 55-57]. Similarly, *MET* and *RET* are among the high-scoring Tclin kinases for Thyroid Cancer (THCA). On average, we find that Tclin kinases had higher CKI scores, suggesting that many of the most clinically relevant or prognostic cancer kinases have already been studied extensively. However, multiple understudied kinases also exhibit a high CKI score thus indicating new likely clinically relevant targets.

**Figure 4:**
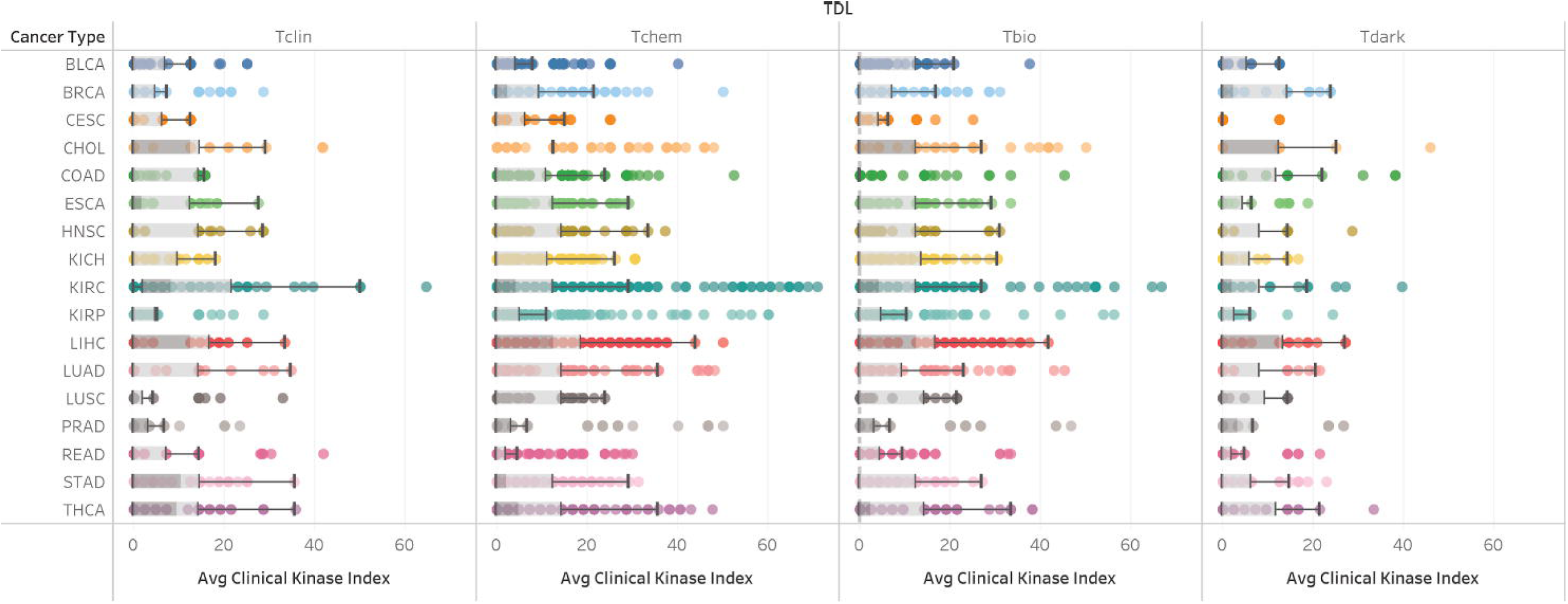
Clinical Kinase Index (CKI) scores by Target Development Level (TDL) and TCGA-Cancer Cohort. There are generally no significant differences in CKI scores amongst TDL with the exception of CHOL, KIRC, and STAD cohorts (Kruskall-Wallis, p<0.05). These cancers have average Tclin scores that are significantly higher than the other target development levels. COAD, BRCA, LUSC and PRAD have Tdark kinases that score on average higher than Tclin kinases.

The highest scoring kinase in all datasets is *PLK1* (Tchem), with a CKI of 70.83 in Kidney Renal Papillary Cancer (KIRP). *PLK1*, like other Tchem kinases, rank very highly, because they are attractive targets that are or have been under pre-clinical or clinical investigation [58, 59]. Data has shown that it is overexpressed in a number of cancers and that its activity has been linked to tumor growth, metastasis and drug resistance. *AURKA* and *AURKB* also consistently scored high across every cancer cohort, as did other cell-cycle and mitoses related kinases (*BUB1B, BUB1, CDK1*). Despite Aurora kinases being priority targets for cancer drug development, clinical trials have, thus far, showed limited efficacy in solid tumors [26, 50].

Understudied kinases rank among the top 20 kinase genes for every cancer cohort scored. Several of these kinases appear to be clinically relevant in multiple cancers. For example, *ERN2* (Tbio) is the 2^nd^ highest scoring kinase in BRCA and CHOL and 6^th^ in LIHC. Due to its understudied nature, there are little to no cancer-related publications available to assess the literature evidence of *ERN2* as a potential target. *ERN2* does code for the protein IRE1β, which is part of the unfolded-protein stress response pathway[60]. The UPR pathway is a pro-survival pathway that is hijacked by cancer cells and thus has been a topic of discussion in the context of drug development [61]. Other understudied kinases that score favorably in the majority of cancer cohorts include *PKMYT1* (Tchem), *DCLK3* (Tchem), *BRSK*1 (Tchem), *ADCK5* (Tdark) and *LMTK3* (Tbio).

To compare the kinase scores across each cancer cohort, we performed a Spearman correlation rank analysis. Correlation between similar cancer cohorts based on tissue seemed to be strongest, with LUSC and LUAD having significant overlap in top ranking kinases, as was the same in KIRC with KIRP and COAD with READ (Figure 5). The top 25% kinases in each cancer were also used as input for MSigDB [62] (Gene Set Enrichment Analysis) to compare gene set enrichment overlap between each cancer. Spearman correlation using rank-ordered gene sets show varying degrees of overlap between all cancer cohorts, but the strongest overlap was between LIHC and CHOL (85 gene sets in common). These data suggest that while many of the same key players are involved in the progression of multiple cancers, unique kinases emerge in each cohort. All cohorts were highly enriched for genes in the “FIRESTEIN_PROLIFERATION” and “MODULE_244” gene sets, which are genes required for the proliferation of colon cancer cells and genes responsible for DNA damage repair, respectively[63].

**Figure 5:**
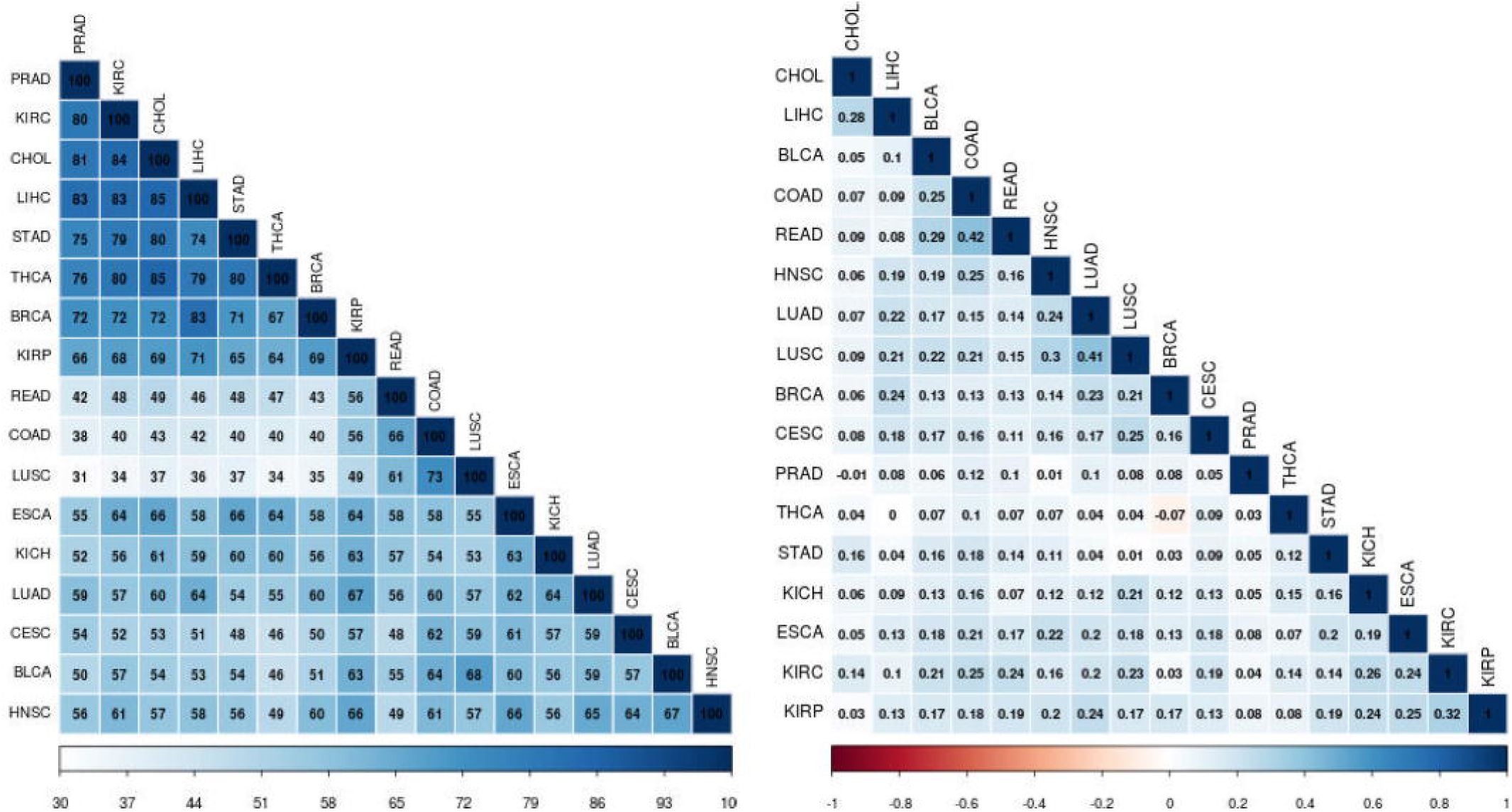
Overlap Analysis between Cancer Cohorts A.) Gene sets enrichment analysis was performed on the top 25% scoring kinases for each cancer cohort using MSigDB. Pairwise overlap of 100 gene sets was calculated. Cohorts which share the highest degree of gene set enrichments include LIHC and CHOL, KIRC and CHOL and THCA and CHOL. B.) Spearman-rank correlation between CKls of each cancer cohort. The greatest degree of correlation was present between cancers of the same tissue, included COAD and READ or LUAD and LUSC. THCA and BRCA had a slight negative correlation, suggestive of opposing clinical relevance of the same kinases in the different cancer cohorts.

### CKI Scores are Supported by ACHILLES Dependency Map (DepMap) Data

Project Achilles and the Dependency Map [64] are efforts to identify genes essential for cancer cell proliferation and survival. Combining RNAi and CRISPR systematic loss of function screens in 563 cancer cell lines for over 17,000 unique genes, researchers were able to obtain gene-level “dependency” scores while accounting for off-target shRNA/Cas-9 effects and other molecular features using the DEMETER computational method [64]. We wanted to evaluate our kinase-ranking predictive model against this experimentally derived data to understand if cancer kinase dependency in cell-models correlated with clinical and pathological features associated with the dysregulation and overexpression of these kinases. Although we would not expect cell line data to perfectly correlate with clinical data, there should be some indication of clinical relevance in these datasets. 565 of the 634 kinase genes in our dataset were present in the Achilles database. Of the 69 kinases that were not present in Achilles, 21 (30%) were understudied, which further underscores the inherent bias against these genes. We first compared the median and average clinical kinase scores between groups of “dependent” (DepMap Score <-1.0) and non-dependent kinases using Kruskall-Wallis analysis. For each cancer type and cell line analyzed (with the exception of THCA), dependent kinases had a significantly higher clinical kinase index and the differences in distributions between the two groups were significant (p<0.05) (Figure 6, Supplemental Table 5). Interestingly, some of the Tclin kinases had high clinical kinase index scores with high dependency scores (not dependent) in each cancer cell line, suggesting cell line data may fail to rank-order these currently approved drug targets. Conversely, some Tclin kinases in our model scored low on the CKI but had significant dependency scores. Further investigation revealed that cyclin-dependent kinases tend to score “low” as mRNA levels do not correspond with activity in tumors. CDKs, as their name suggest, have activity that is dependent on the abundance of cyclins. Also, as previously mentioned, kinases such as mTOR have activity that is related to its post-translational modification. Despite these discrepancies, we can make the case that our kinase target ranking based off of clinical phenotypes correlates to a degree with cancer cell dependency in validated cancer cellular models. Receiver Operator Characteristic Curves (ROC) were also generated to compare our predictive model against DepMap Data. Using a cutoff of <-1 as “good”, we find that the average CKI (across all cancers) is predictive of average DepMap score (across all cancer cell lines) with an ROC score of 0.776 (Figure 6).

**Figure 6:**
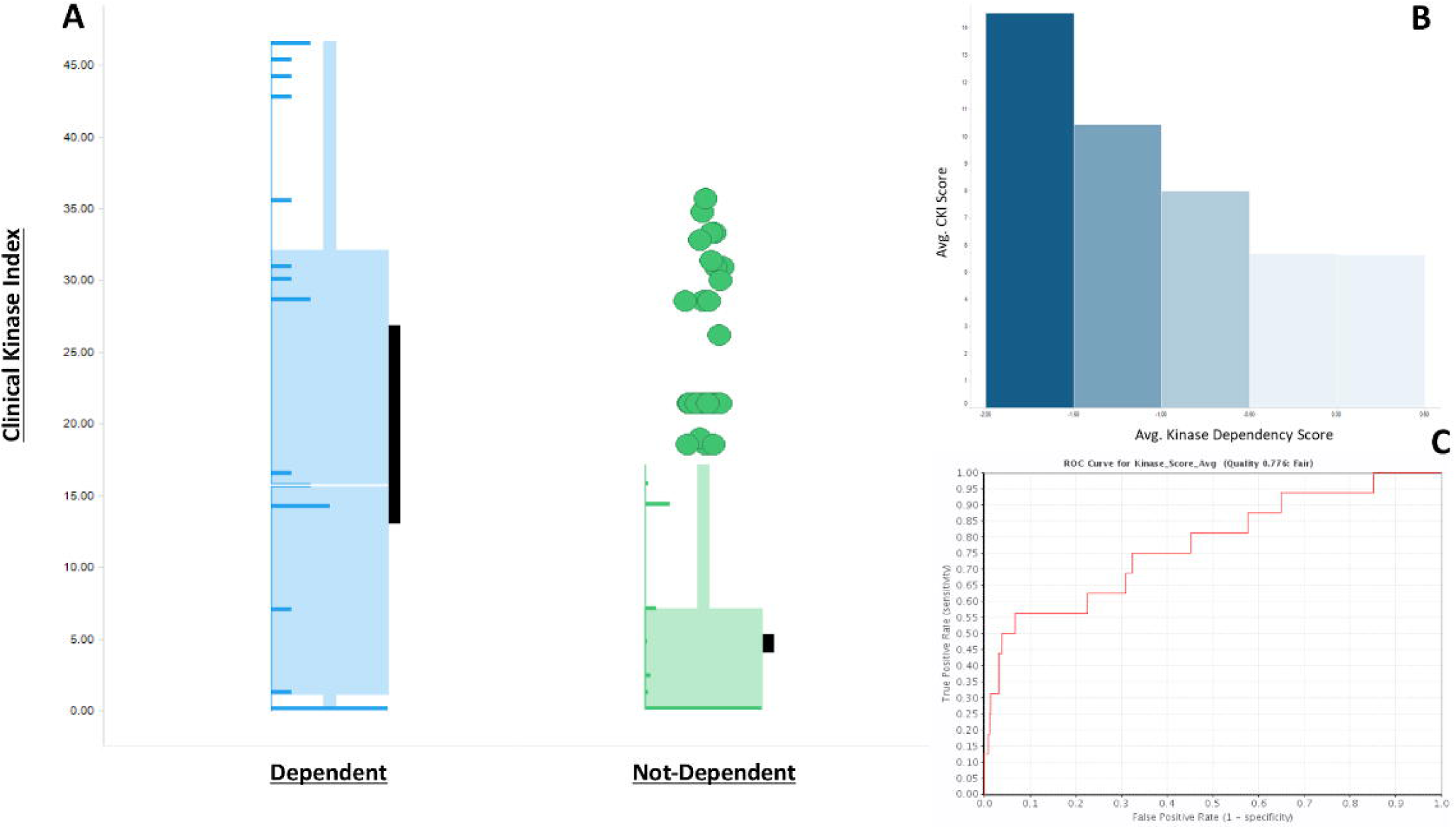
Comparison of Dependency Map Scores and CKI. A.) Kinases from the ACH-000851 cell line (lung cancer) were divided into “Dependent” (DepMap Score <-1.0) and “Not-Dependent” Groups. LUSC Clinical Kinase Index scores were compared between the groups using the Kruskall-Wallis test. Kinases that are dependent in lung cancer cell lines also score significantly higher in the clinical kinase index (p=2.13E-07). Black bars represent the 95% confidence interval of the mean. Distribution of scores is depicted as horizontal bar graphs superimposed on the box-plot. B.) As DepMap Scores (Binned) increase (becoming less dependent), CKI scores decrease. C.) ROC curve data using average CKI score and average Dependency score with a cutoff of <-1.0 shows that DepMap data is predictive of clinical relevance (ROC=0.776).

## Discussion

In this study, we systematically integrated differential gene expression (DGE), Kaplan-Meier survival, mutational hot-spot and clinical/pathological correlation analyses in order to prioritize clinically relevant kinase targets across 19 TCGA cancer types. Our clinically focused pan-cancer and pan-kinome analysis highlighted multiple understudied kinases with the potential of promising druggable target opportunities. Moreover, through the development of our accompanying Clinical Kinase Index App, which is freely available at http://cki.ccs.miami.edu, we facilitated the discovery, exploration and analysis of our data by the scientific community. Our comprehensive analysis across 19 cancer types and our plethora of rich metadata annotations (e.g. Target Development Level, approved MOA, kinase phylogenetic and functional classifications), offer researchers a quick and intuitive way to explore the entire kinome and identify understudied kinases with a high therapeutic potential. This app offers an intuitive and comprehensive view of the kinome based on cancer type-specific differential expression, survival data, and TNM pathological staging. Researchers can obtain a prioritized list of dark and understudied kinases based on multiple criteria (e.g. cancer type, kinase family). Our rich metadata annotations enable researchers to explore our data via Target Development level (TDL), approved Mechanism of Action (MOA), and kinase classifications (including pseudokinases, non-protein kinases etc.). This unique capability can drive a more efficient drug target prioritization by the research communities. For each dark / understudied kinase, researchers can obtain cancer type-specific analysis and rank-ordered prioritizations. As more information is discovered and more bioinformatics tools and workflows are available such as phosphoproteomics and active kinome profiling, the CKI will be updated and optimized to continually enrich the dataset for clinically relevant kinase targets.

Many researchers have combed the TCGA to find prognostic genes for cancer. However, none have focused their efforts on the kinome, in particular, the understudied kinome. While the TCGA has multi-dimensional data (proteomic, genomic & transcriptomic), it has been suggested that mRNA gene expression alone has the best prognostic power compared to mutational, proteomic or methylation data [65]. Gene expression data is also the most abundant in many public data repositories. The CKI therefore has been primarily based on the significance of gene changes with respect to various clinical parameters. Several kinases discussed in this study and present in our analysis are likely to be important drug targets for cancer.

Comparing the average CKI scores of “Understudied” vs. “Studied” kinases across all cancers and within individual cancer cohorts reveals no significant difference (except for in the CESC Cohort, p=0.0083), suggesting that there are many understudied kinases with clinical relevance comparable to currently approved kinase drug targets. The highest scoring understudied kinase is PKMYT1. PKMYT1 is a member of the WEE1 family of kinases that negatively regulates the G2/M transition of the cell cycle by phosphorylating and inactivating cyclin-dependent kinase 1 [66]. PKMYT1 is overexpressed in 17 of the 20 cancers analyzed in this study. Cox-regression analysis of PKMYT1 reveals this kinase is a powerful prognostic and predictive biomarker for survival in KIRC and KIRP cohorts. Other computational work has identified PKMYT1 is a novel drug target for kidney cancer, using co-expression analysis to reveal PKMYT1 clusters with other important cell cycle genes[67]. PKMYT1 is classified as Tchem with several active compounds, including PD-0166285, IC50 = 7 nM and PD-173955, Kd = 44 nM. Although these compounds are not considered chemical probes, they provide starting points for the development of a selective probe or a viable lead compound.

The expression of many understudied kinases is believed to correlate with the clinical progression of several cancers. RIOK1 (Tchem) is one such kinase whose mRNA levels are linked to metastasis in 4 cancers (BLCA, KICH, KIRP, LUAD). Recent studies have identified RIOK1 as a novel drug target and dependency map data shows RIOK1 as a common highly essential gene in 561/563 cell lines tested. In fact, data from ChEMBL lists sunitinib has been reported as s a known kinase inhibitor of RIOK1 with a Ki of < 35nM. However, more selective inhibitors of RIOK1 will be necessary to further validate it as a therapeutic target of interest. We have also identified 67 other understudied kinases that have mRNA levels prognostic of survival and progression in at least two cancer cohorts. All Kaplan-Meier survival curves and corresponding p-values are available for viewing via our Clinical Kinase Index (CKI) Web application.

While many of the high scoring novel kinases are also beginning to be discussed in other papers, many are too “dark” and thus warrant more exploration. Even in large scale datasets and studies, dark kinases are excluded from the analyses due to lack of validated antibodies, assay, and chemical probes and lack of interest and knowledge of the biology of these targets. A large-scale concerted effort must be taken to effectively bring these “dark” kinases into the light. Such target validation efforts must include RNAi/CRISPR knockdown/knockout cell lines and mice models, elucidation of co-expression and regulatory networks, substrate identification and assay development, and the development of highly specific chemical probes and the determination of co-crystal structures to facilitate the optimization of lead compounds for future drug development efforts. Much of this work is currently pursued in Illuminating the druggable Genome (IDG) program (https://druggablegenome.net/). The results of this paper will help us, and other groups, select and prioritize their target of choice based off of the clinical and biological focus of their own research interests. 2018 was a record year for FDA approvals, yet only 3 of the 39 drugs targeted novel kinase targets and moved target development levels from Tchem to Tclin [68]. The rate at which kinase targets move from Tbio or Tdark (little known biology and no potent chemical probes) to Tchem (a small molecule probe with Kd <30nm) is alarmingly slow. No more than 20 IDG targets move from Tbio to Tchem each year, a number encompassing *all* target types (kinases, GPCRs, ion channels). By this pace, it would take decades to wholly illuminate the kinome. Thus, the biases against understudied targets must be lifted particularly in grant funding so that there can a full-fledged concerted effort to explore the druggable genome.

## *STAR* Methods

### Inclusion Criteria: TCGA Datasets

The TCGA data portal contains the molecular data of over 20,000 tumor and matched normal samples for 33 cancer types from over 11,000 patients (https://www.cancer.gov/tcga). Our inclusion criteria for kinase analyses were based on the availability of data including normal samples available for differential gene expression analysis and mutational hotspot analysis, clinical data and survival data. First, we only considered solid tumors, thus excluding LAML and DLBC. 9 Cancers (OV, THYM, UVM, SARC, PCPG, UCEC, GBM, UCS and LGG) have been excluded from the scoring system due to lack of clinical data (TNM staging). 5 Cancers have only clinical and survival data (ACC, SKCM, PAAD, MESO, and TGCT) but not enough normal samples to perform a differential gene expression analysis or mutational analysis. 17 Cancers had the full requirements for scoring for this paper, but all data and analyses are available via our Clinical Kinase Index (CKI) application. An overview of the TCGA dataset with number of normal samples, tumor samples and data availability is compiled in the supplementary material (Supplemental table 6). Abbreviations for all the cancer types used in this paper are in accord with the TCGA data portal.

### Inclusion Criteria: Kinases

The complete list of kinases used in this analysis was obtained from Pharos (https://pharos.nih.gov/) [23]. 634 kinases were included in our initial analyses; this list contained protein kinases, non-protein kinases and pseudokinases. Several of the kinases annotated in Pharos have different gene names in the TCGA dataset, and needed to be manually curated for our data. Kinase names that could not be mapped to Pharos or whose expression levels were undetectable after expression normalization were excluded from our analyses. Target development levels (TDL) were also obtained from Pharos. Kinases were additionally grouped as “Understudied” or Studied, based off of the IDG designation also available from Pharos. As of June 2019, 151 kinases were annotated as Understudied. However, only 149 of these were included in our analyses (MAP3K21 and STK19 were excluded for above reasons).

Kinases were finally annotated, per cancer, as MOA targets of approved drugs indicated for the treatment of that specific cancer. For example, while BRAF may have a Tclin annotation, it is not a primary clinical drug target for Breast Cancer (BRCA) treatment. Data for drug target MOA and drug indication was obtained from DrugCentral (http://drugcentral.org/) [1, 43] and “A Comprehensive Map of Molecular Targets” [69] (Supplemental table 7). BLCA was modified to include FGFR1/2/3/4 due to the 2019 approval of Erdafitinib [70]. Kinase group and family information were obtained from Kinase Drug Target Ontology (DTO) database [11].

### Data extraction and preprocessing

The TCGA data were downloaded from recount2[71] where TCGA clinical data are also available for all 31 cancers. Differential expression analysis was performed for 19 of the 31 cancer types. Datasets with at least three normal (control) samples and three tumor samples were considered, and all other datasets were discarded. For the 12 cancers (ACC, LGG, SARC, DLBC, THYM, OV, SKCM, LAML, UVM, PAAD, TGCT, UCS) data were not available for normal samples or were below three normal samples thus preventing the comparison between normal and tumor samples. The data downloaded from recount2 was pre-processed by removing the duplicated TCGA patient IDs or barcodes in each cancer type. Further, the counts were scaled and the lowly expressed genes filtered out using R limma package[72] The resultant datasets for each cancer were used for differential expression (DE) analysis, survival analysis, identification of mutational hotspots, and analysis of clinical significance (TNM, Path, Histology).

### Clinical Data

Patient data for AJCC Pathological Stage[73] and histological grade for each cancer type was obtained from the standardized clinical dataset, the TCGA Pan-Cancer Clinical Data Resource[74] T,N, and M Staging was obtained from the GDC [75] database using the R/Bioconductor[76] package recount2[71]. Unique patient IDs were matched using R for each cancer type to include only primary tumor samples for clinical analyses (Sample code “01”). Secondary or metastatic tumors were not considered in this analysis.

### Differential Gene Expression Analysis

For the differential expression analysis, we applied the TCGAbiolinks pipeline on the filtered genes for 19 cancers[77]. Genes with log_2_FC > 1 and FDR threshold of 0.01 were considered significantly DE. The significantly DE genes were mapped to the kinases per cancer. Specifically, only the overexpressed kinases from each cancer type were further considered for the scoring analysis.

### Survival Analysis

In addition to gene expression analysis across cancer samples, survival analysis based on gene expression levels was performed. Available TCGA patient data were used to generate Kaplan-Meier (KM) survival plots. For the plots, patient clinical data was obtained with, i) patient vital status (Alive or Dead), ii) time (if patient is alive then, “days_to_last_follow_up”, if patient is dead then, “days_to_death”), and iii) expression level. Patients with no vital status or follow_up data were considered censored.

For each kinase, the Kaplan-Meier (KM) survival plot was generated across each cancer type, applying the survival R package[78] The patients were categorized into two distinct groups, high expression (upper quartile) and low expression (lower quartile). The resulted KM survival curves were compared by log-rank test obtaining a P-value, which indicates statistical significance of survival samples. Kinases with high expression and significant P-value were used for score analysis.

### Mutational Hotspot Analysis

Mutation somatic variants data was obtained from the GDC data portal in the Mutation Annotation Format (MAF) for each cancer type[38]. The pre-compiled TCGA MAF objects including somatic mutation along with clinical information were downloaded from the TCGAmutation R package[76]. All Kinases were mapped to each cancer type using R. In addition, the function oncodrive was applied to identify cancer genes (kinase drivers) from a given MAF file[38]. Oncodrive is a function based on oncodriveCLUST algorithm, which has been implemented in Python[79]. For scoring analysis, specifically, only those kinases were selected for each cancer that had a minimum of 5 mutations and significant P-values. Kaplan-Meier curves for mutated or amplified kinases were generated using cBioPortal[80].

### Clinical Analysis

For clinical Analysis, Tumor, Node, Metastasis (TNM) staging system developed by American Joint Committee on Cancer AAJC[73]was used. The TNM system classifies cancer:

1. T: by the size, which is further grouped into t1- (t1a,t1a1,t1b, t1b1), t2- (t2a,t2a1,t2a2,t2b,t2c), t3- (t3a,t3b,t3c) and t4- (t4a,t4b,t4c,t4d, t4e).
2. N: involvement of regional lymph nodes, subtypes n0, n1- (n1a, n1b, n1c, n1mi), n2- (n2a, n2c, n3a) and n3 - (n3a,n3b,n3c).
3. M: presence or absence of distant metastasis, sub divided into m0 and m1.

We used ANOVA to identify significant differences between t1-t4 for each of the four metrics. Subsequently, ANOVA was also performed for N and M to see if there are any significant changes between n0-n3 and m0-m1. In addition, the histological grade was also considered for scoring. The grade is a qualitative assessment for differentiated cells under microscope. The differentiated cells are low grade (g1, g2) and dysmorphic and de-differentiated cells are considered high grade (g3, g4). The statistical significance differences between histological grade g1-g4 was calculated using ANOVA.

### Scoring

We have developed the Clinical Kinome Index (CKI) to evaluate the prognostic and clinical value of the mRNA expression levels of each kinase in the human kinome. RNASeq data obtained from normal and solid-tumor samples was used to correlate RPKM with advanced tumor staging in clinical, pathological and histological classifications. Since we are mostly interested in the understudied kinome, we opted to not include RPPA or protein expression data since this was unavailable due to lack of specific and validated antibodies for such kinases. Additionally, we recognize that levels of messenger RNA do not necessarily correlate linearly with kinase activity[22]and a future study of the phospho-kinome would offer more insight into effects these kinases have on cancer progression.

The Clinical Kinome Index (CKI) score was generated for each kinase per cancer using 4 parameters: Differential Gene Over-Expression, Overall Survival (OS), Mutational Hotspots and Clinical/Pathological Progression. If the kinase was shown to be significantly overexpressed, correlated with negative survival outcomes or significantly mutated, it received a score of 1 (per parameter). Clinical scores were created using an average per each clinical parameter which include clinical stage, T, N and M staging, and histological staging. For example, ANOVA analysis was used to determine significant differences in the means of expression for each kinase in each T stage (t1 vs t2, t2 vs t3 etc.). If a kinase expression was significantly increased between two T stages, it received a score of 1. If it was significant between multiple T stages, the scores were averaged to have a maximum score of 1. This was repeated for M and N stages as well as other clinical parameters including histological grade, pathological stage and clinical stage. The total clinical score was a sum of these scores, with a maximum score varying per cancer due the differing availability of clinical and pathological data in the TCGA datasets. Final scores were determined by summing all parameters and dividing by the maximum possible score and multiplying by 100% to normalize across all cancer cohorts. In short, a score is only assigned to a kinase if its increased expression correlates with a progression in staging, a decrease in survival or has significant mutations.

### Clinical Kinase Index (CKI) Web Application

The Clinical Kinase Index and application was developed and written in R language R 3.3 or higher using the R packages: shiny, RColorBrewer, gplots, plyr, ggplot2, limma, TCGAbiolinks, tidyverse, recount2, edgeR, SummarizedExperiment, devtools, superheat, xlsx, survival, RTCGA.rnaseq,, RTCGA.clinical, survminer, maftools, corrplot, colorRamp, and plotly. The construction of the App utilizes Shiny, a framework to build and deploy interactive Web applications in R. The data have been processed using the R Bioconductor databases and packages included within other requirements. The CKI homepage includes the mRNA expression heatmaps of all kinases (and understudied kinases). The application includes a table which can be filtered based off of user input and contains the kinase gene name, cancer of interest, clinical kinase index score, ranking and other meta-data annotations for the target. Through the gene page users can explore the score of a particular kinases of interest taking advantage of all the information available for specific dataset, which can be filtered: TDL, Kinase type, Kinase Group, MOA targets. Additionally, the CKI disease page allows users to generate different plots of particular cancer and gene of interest including 1) Volcano plots to display differentially expressed kinases per cancer, 2) Boxplots showing the expression of kinases according to TNM staging, and 3) Kaplan Meier curve representing the overall survival levels of each gene per cancer. The data to reproduce the plots can be downloaded on the Download page for future analysis. This includes some data that has not been visualized in the app, such as mutational and CNA analyses.

### DepMap Analysis

Data was downloaded from DepMap [64]. The datasets used were CRISPR (Avana) Public 19Q2 and the cell line metadata. Using DepMap Cell line metadata, cell lines were extracted using “disease” and “disease subtype” terms to match the correct TCGA dataset.

### External Tools

Gene amplification data was obtained from cBioPortal using all TCGA Provisional Datasets. Focal amplifications were identified using the GISTIC tool from the BROAD Institute[44]. Gene Set Enrichment Analysis was performed using MSigDB[62]. The top 25% scoring kinases for each cancer were used as input. Positional Gene Sets, chemical and genetic perturbations, canonical pathways, KEGG gene sets, cancer modules and oncogenic signatures were selected for analysis. The FDR q-value threshold was 0.05 and the top 100 enriched gene sets were saved.

## Author Contributions

D.E., V.S., and S.C.S. conceived the project and R.K. carried out statistical analyses. D.E. analyzed and interpreted data and R.K. developed the Web app. D.E. wrote the manuscript and D.E., V.S. and R.K. made figures.

## Acknowledgments

This work was supported by NIH grants U54HL127624 (BD2K LINCS Data Coordination and Integration Center, DCIC), U24TR002278 (Illuminating the Druggable Genome Resource Dissemination and Outreach Center, IDG-RDOC), and U01LM012630 (BD2K, Enhancing the efficiency and effectiveness of digital curation for biomedical ‘big data’), and the State of Florida Biomedical Research Program, Bankhead Coley grant number 9BC13.

