## Supplemental Figures for "The Clinical Kinase Index: Prioritizing Understudied Kinases as Targets for the Treatment of Cancer"

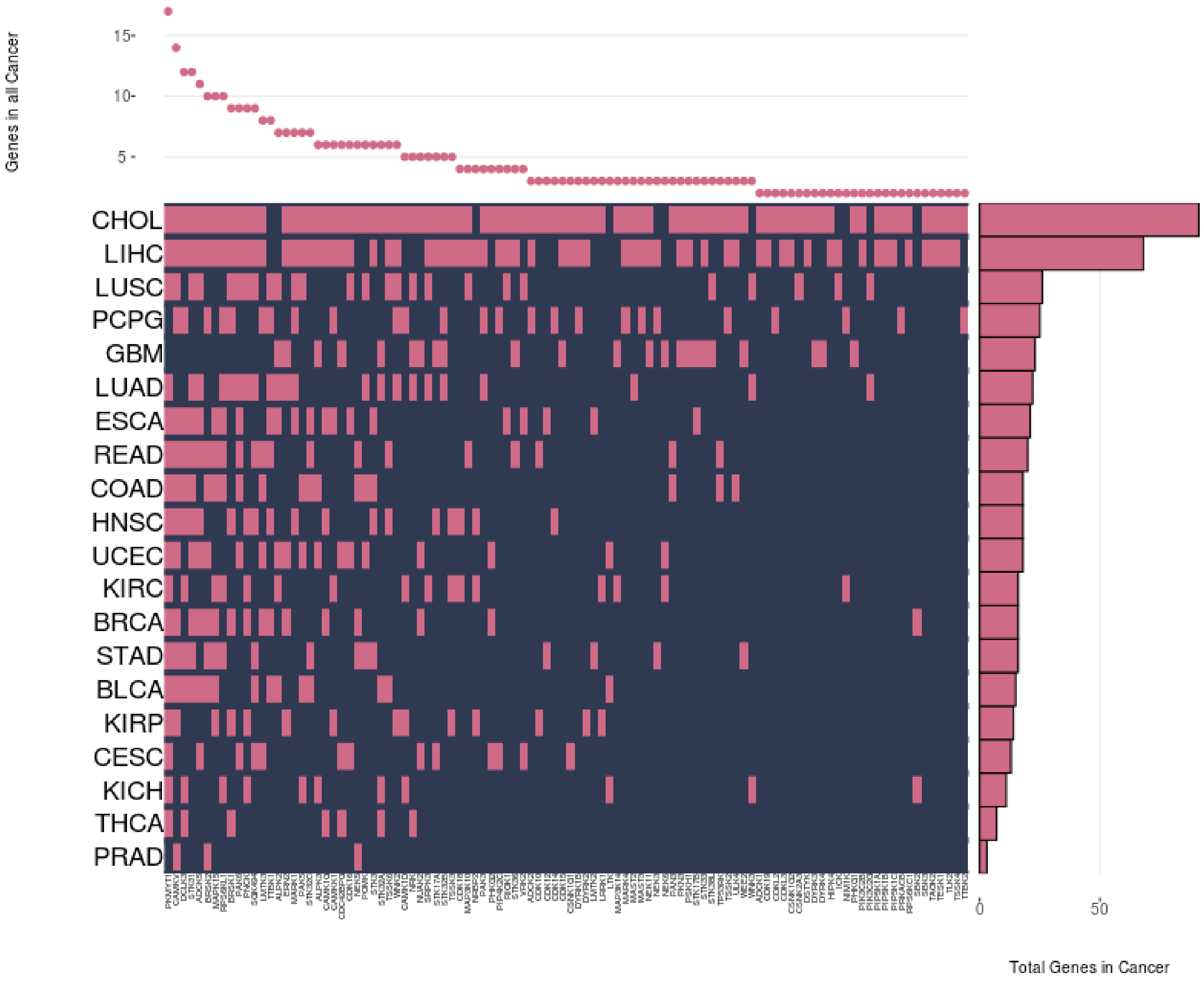


Figure S1: Differential overexpression of Understudied kinases in each cancer cohort. 102 Understudied are significantly overexpressed in at-least two cancers.

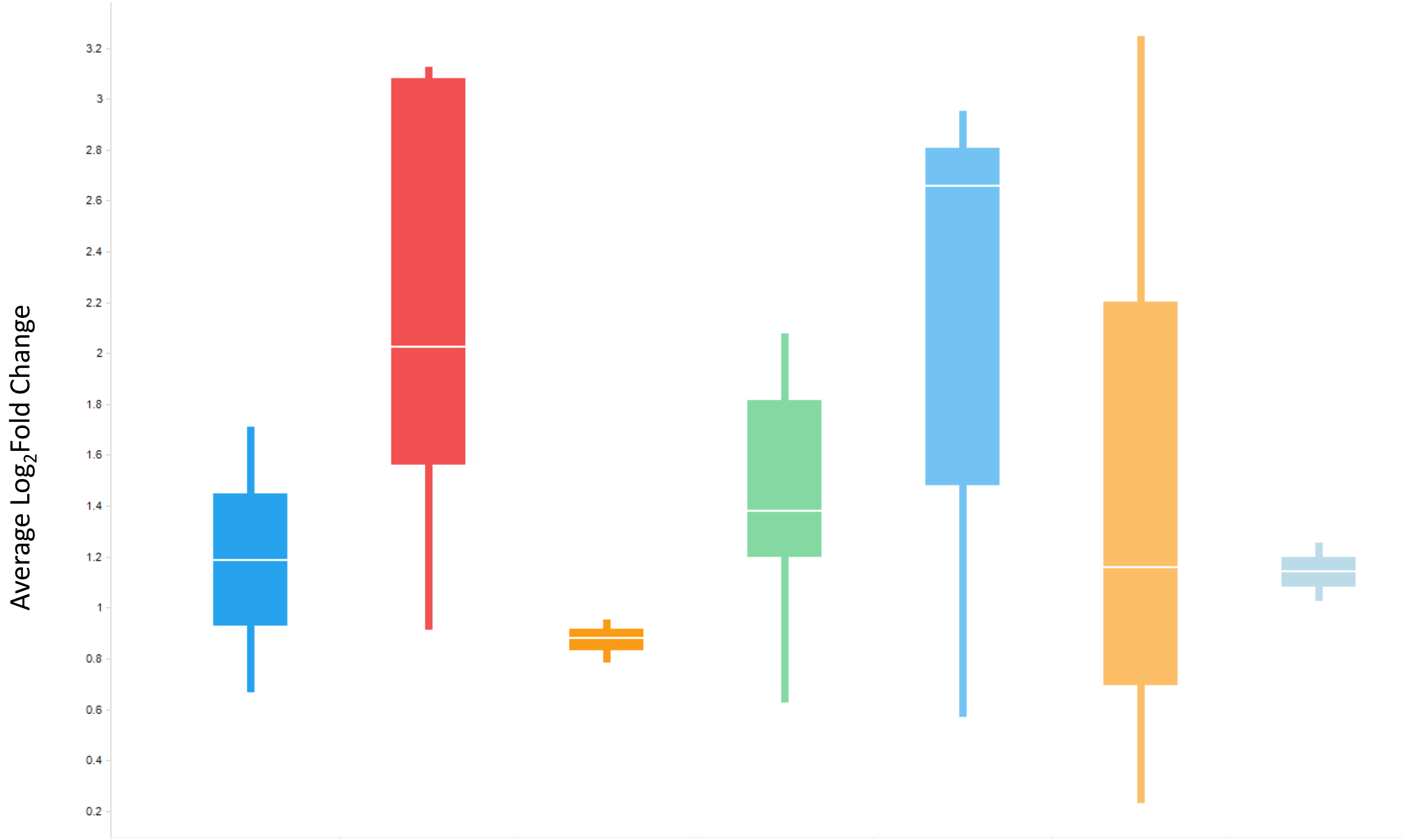


| | AGC | CAMK | CMGC | Nucleoside/Nucleotide | Other | TK | TKL |
|---|---|---|---|---|---|---|---|
| Median | 1.18819 | 2.02485 | .88034 | 1.38002 | 2.65938 | 1.15854 | 1.14054 |
| Avg | 1.18819 | 2.14169 | .871481 | 1.42711 | 2.14167 | 1.54538 | 1.4054 |

Figure S2: Several kinase groups appear to be enriched in their overexpression in various cancers. Averaging the log2FC of all kinases across all cancers demonstrates that certain kinase transcripts are consistently very highly upregulated. Kinases from the understudied CAMK group have the highest average log2FC (2.14) and kinases from the Other group have the highest median log2FC (2.66).

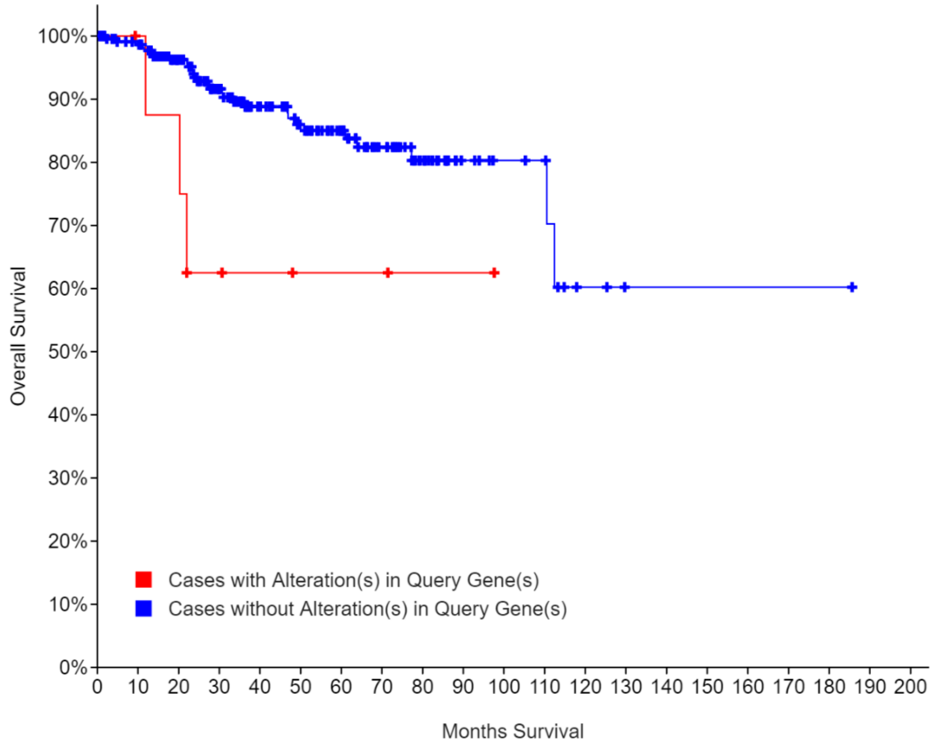


Figure S3: *DYRK1B* is mutated in 4.27% of Endometrial Carcinoma samples (UCEC) and Kaplan-meier analaysis shows that those with mutations in this kinase have worse overall survival (OS) (p-Value= .0204).

Figure S4: *Tdark* Kinases are potential prognostic biomarkers for several cancers. A.) Tdark Kinases which are significant for survival. ALPK3 has mRNA levels which are prognostic for survival in 6 cancer cohorts. B).TNM staging scores (0-3) show that dark kinases contribute to tumor progression.

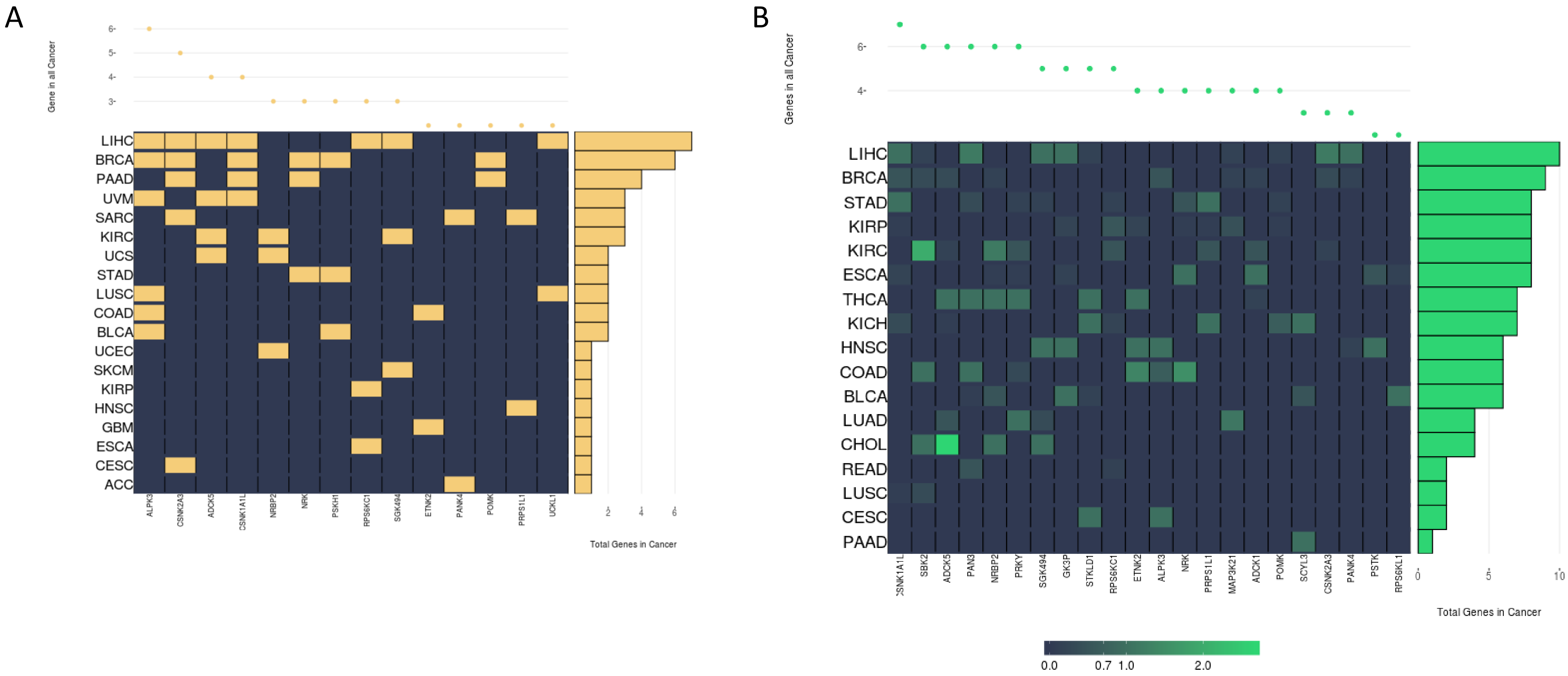
